# Improved antibody structure prediction by deep learning of side chain conformations

**DOI:** 10.1101/2021.09.22.461349

**Authors:** Deniz Akpinaroglu, Jeffrey A. Ruffolo, Sai Pooja Mahajan, Jeffrey J. Gray

## Abstract

Antibody engineering is becoming increasingly popular in medicine for the development of diagnostics and immunotherapies. Antibody function relies largely on the recognition and binding of antigenic epitopes via the loops in the complementarity determining regions. Hence, accurate high-resolution modeling of these loops is essential for effective antibody engineering and design. Deep learning methods have previously been shown to effectively predict antibody backbone structures described as a set of inter-residue distances and orientations. However, antigen binding is also dependent on the specific conformations of surface side chains. To address this shortcoming, we created DeepSCAb: a deep learning method that predicts inter-residue geometries as well as side chain dihedrals of the antibody variable fragment. The network requires only sequence as input, rendering it particularly useful for antibodies without any known backbone conformations. Rotamer predictions use an interpretable self-attention layer, which learns to identify structurally conserved anchor positions across several species. We evaluate the performance of our model for discriminating near-native structures from sets of decoys and find that DeepSCAb outperforms similar methods lacking side chain context. When compared to alternative rotamer repacking methods, which require an input backbone structure, DeepSCAb predicts side chain conformations competitively. Our findings suggest that DeepSCAb improves antibody structure prediction with accurate side chain modeling and is adaptable to applications in docking of antibody-antigen complexes and design of new therapeutic antibody sequences.

## Introduction

Antibodies are specialized proteins that play a crucial role in the detection and destruction of pathogens. The binding and specificity of antibodies are largely determined by the complementarity determining regions (CDRs) that consist of three loops in the light chain and three loops in the heavy chain [1]. Structural diversity is largely achieved by the third loop in the heavy chain, which determines many antigen binding properties. Additionally, CDR H3 does not have a canonical fold like the other loops [2], making it challenging to model [3–5]. Currently, engineering of new antibodies is hindered by accurate prediction of the CDR H3 loop, including the corresponding side chains for docking applications. Prediction of side chains is a critical component of structure prediction and protein design [6]. The surface of the antibody CDR loops including the side chains play an important role in antigen recognition [7].

There has been a growing interest in effective design of new antibodies since they are commonly used in biotherapeutics [8]. Antibody structure determination via techniques like X-ray crystallography and NMR is challenging and time-consuming. Machine learning methods improve overall structure prediction and docking [9]. Recently, highly accurate structure prediction models have been proposed for proteins in general [10–12] and for antibodies [13–16]. The performance of AlphaFold2 has been impressively accurate in the recent CASP14 experiment and surpassed most other protein structure prediction methods proposed to date [11]. Unlike the other deep learning-based methods, AlphaFold2 predicts all side chain rotamers in addition to the protein backbone. While current antibody prediction methods utilizing deep learning do not directly predict side chains, they are all able to predict the backbones with high accuracy. Hence, a next step towards the advancement of antibody modeling and engineering is the accurate prediction of side chains to improve overall structure prediction and docking.

Presently, there are successful methods for rotamer predictions that rely on calculating the probability of a *χ* angle as a function of backbone torsion angles. For instance, SCWRL4 uses backbone-dependent libraries to calculate rotamer frequencies based on kernel density estimates and kernel regressions [17]. Antibody-specific methods like PEARS capture rotameric preferences based on the immunogenetics numbering scheme to restrict possible side chain conformations in the sample space based on positional information [18]. Both SCWRL4 and PEARS require the antibody sequence and backbone structure to generate side chain predictions. They repack the side chains onto the provided backbone, and their performance generally declines when the input is not the crystal backbone. To address these limitations, we propose DeepSCAb (deep side-chain antibody), a deep neural network that predicts full *F*_V_ structures, including side chain conformations from only the amino acid sequence.

## Methods

### Antibody structure datasets

#### Training dataset

The training dataset for our model was curated from SAbDab, a database of antibody structures from the Protein Data Bank [19]. We enforced a threshold of 99% sequence identity as well as a resolution cutoff of 3 Å for high quality data. We removed targets belonging to the RosettaAntibody benchmark set [20] to evaluate model performance, which resulted in a total of 1433 antibody structures that we used to train and validate our network.

### Predicting antibody structure from sequence

DeepSCAb consists of two main components: an inter-residue module for predicting backbone geometries and a rotamer module for predicting side chain dihedrals. The inter-residue module is initially trained separately and then in parallel with the rotamer module.

#### Simultaneous prediction of side chain and backbone geometries

The initial layers of the model for predicting pairwise distances and orientations are based on a network architecture similar to that of DeepH3 [13]. The inter-residue module consists of a 3 block 1D ResNet and a 25 block 2D ResNet. The one-hot encoded antibody sequence of dimension *L* × 21 is taken as input to pass through the module via the 1D ResNet, where *L* represents the number of residues in the sequence and 21 represents each amino acid type with the addition of a chain delimiter. The 1D ResNet begins with a 1D convolution that projects the input features up to *L* × 64, followed by three 1D ResNet blocks (two 1D convolution with a kernel size of 17) that maintain dimensionality. The output of the 1D ResNet is then transformed to pairwise by redundantly expanding the *L* × 32 tensor to an *L* × *L* × 64 tensor. Next, this tensor passes through 25 blocks in the 2D ResNet that maintain dimensionality with two 2D convolutions and kernel size of 5 × 5. The resulting tensors are converted to pairwise probability distributions over *C_β_* distance, *d*, the orientation dihedrals *ω* and *θ*, and the planar angle *ϕ*. The inter-residue module is trained as described for DeepH3 [13].

The rotamer module takes as input the inter-residue features. The tensors of dimension *L* × *L* × 64 resulting from the 2D ResNet are transformed to sequential by stacking of rows and columns to obtain a final dimension of *L* × 128. The rotamer module contains a multi-head attention layer of 1 block with 8 parallel attention heads and a feedforward dimension of 512. The self-attention layer outputs *L* × 128 tensors, which then pass through a 1D convolution with kernel size of 5. The tensors are converted to rotamer probability distributions that are conditionally predicted for each *χ* dihedral using softmax. For example, *χ*_1_ is an input to *χ*_2_, and *χ*_1_ through *χ*_4_ are inputs to *χ*_5_. The predicted rotamers are added back into the inter-residue module: the rotamer tensors are stacked onto the pairwise before the final 2D convolution to update the *d*, *ω*, *θ*, and *ϕ* outputs.

Distances are discretized into 36 equal-sized bins in the range of 0 to 18 Å. All dihedral outputs of the network are discretized into 36 equal-sized bins in the range of −180° to 180° with the exception of *χ*_1_. The *χ*_1_ dihedral is discretized into 36 non-uniform bins, with 6 bins of 30° and 30 bins of 6°. The small bins are centered around −60°, 60°, and 180°, consistent with observed conformational isomers. The planar angle *ϕ* is discretized into 36 equal-sized bins with range 0 to 180°. Pairwise dihedrals are not calculated for glycine residues due to the absence of a *C_β_* atom. Side chain dihedrals were not calculated for glycine and alanine residues due to the absence of a *C_γ_* atom and for proline residues due to its non-rotameric nature.

Categorical cross-entropy loss is calculated for each output, where the pairwise losses are summed with equal weight and the rotamer losses are scaled based on each dihedral’s frequency of observation: i.e., *χ*_5_ rotamers are much less frequent than *χ*_1_. The Adam optimizer is used with a learning rate of 0.001. We trained five models on random 95/5 training/validation splits and averaged over model predictions to generate potentials for downstream applications. DeepSCAb models were trained on one NVIDIA K80 GPU, which required approximately 100 hours for 120 epochs of training.

#### Side chain only predictions

To investigate the effect of inter-residual predictions on rotameric predictions, we designed a side chain only network as a control. The control network takes as input the one-hot encoded antibody sequence, which passes through a 3 block 1D ResNet. The remaining architecture of the control network as well as its training process is similar to the rotamer module of DeepSCAb (SFigure 1). However, there are differences in dimension due to the 1D ResNet returning an *L* × 32 tensor. The control network models were trained on one NVIDIA K80 GPU, which required 10 hours for 20 epochs of training. We adopted a shorter training process for the control network as the models tended to overfit after 20 epochs.

### Self-attention implementation and interpretation

#### Transformer encoder attention layer

The rotamer module contains a transformer encoder layer that adds the capacity to aggregate information over the entire sequence (SFigure 2). We tuned the number of parallel attention heads, the feedforward dimension, and the number of blocks according to validation loss during training. We found that 8 attention heads outperformed 16, feedforward with a dimension of 512 outperformed 1,024 and 2,048, and one block of attention performed identically to two. We further experimented with adding a sinusoidal positional embedding prior to the self-attention layer and obtained identical results, implying that the convolutions in our network contain sufficient information on the order of input elements, rendering positional encoding unnecessary [21].

#### Interpreting the attention layer

In our interpretation of the rotamer attention, we take into consideration only one model out of the five that were trained on random training/validation splits. We do not report an average over the attention matrices from multiple models since they vary amongst themselves (SFigure 3). Nevertheless, the properties of attention are conserved across individual models.

We utilize a selected subset of the independent test set to display the most variation across the highly-attended positions as well as the corresponding residue types. This subset consists of the following human PDBs: 1JFQ, 1MFA, 2VXV, 3E8U, 3GIZ, 3HC4, 3LIZ, 3MXW, 3OZ9, and 4NZU.

### Modeling side chains with DeepSCAb in Rosetta

DeepSCAb generates constraints that are utilized for the prediction of an antibody structure. Discrete potentials are converted to continuous function via the built-in Rosetta spline function. The constraints include all 9 geometries, namely *d*, *ω*, *θ*, *ϕ*, *χ*_1_, *χ*_2_, *χ*_3_, *χ*_4_, and *χ*_5_. The *ConstraintSetMover* in Rosetta applies these constraints onto the native pose and then the *PackRotamersMover* models side chain structures. We chose the standard *ref2015* full-atom score function with a weight of 1.0 for all constraints. This protocol can repack side chains on any backbone structure with DeepSCAb predictions.

### Side chain predictions using alternative methods

To assess the side chain prediction accuracy against relative solvent accessible surface area, we compare DeepSCAb to three alternative methods: PEARS, SCWRL4, and Rosetta. For each alternative method, we provide the backbones of the benchmark targets and their sequences. PEARS utilizes antibody-specific rotamer libraries and assigns rotamers based on the IMGT numbering scheme. We generated predictions from PEARS using the publicly available server [18]. SCWRL4 generates *χ* kernel density estimates based on backbone-dependent rotamer libraries by minimizing the conformational energies for each residue. We generated SCWRL4 predictions using the SCWRL4.0 algorithm [17]. Rosetta predictions were generated using default packing protocol (*PackRotamersMover*) and the *ref2015* energy function.

### Data and Code Availability

The source code for DeepSCAb, as well as pretrained models, will be made available prior to publication. The structures predicted by DeepSCAb and alternative methods for benchmarking will be made available prior to publication.

## Results

### Overview of the method

Our deep learning method for antibody structure prediction consists of inter-residue and rotamer modules. We trained DeepSCAb to predict antibody backbones as inter-residual distance and orientations. Then, we simultaneously trained the model to predict the side chain conformations using an attention layer. The pairwise and rotamer probability distributions from DeepSCAb predictions were used for structure realization and packing of the side chains using Rosetta.

### DeepSCAb predicts inter-residue and side chain orientations from sequence

DeepSCAb is a neural network that only requires an antibody sequence to predict full *F*_V_ structures including side chain geometries (Figure 1A). The combined sequences of the antibody heavy and light chains are inputted as a one-hot encoding to initially pass through the inter-residue module. The architecture of this module is similar to our previous method for CDR H3 loop structure prediction, DeepH3 (Figure 1B) [13]. The network is pretrained to predict pairwise geometries, such as *d*, *ω*, *θ*, and *ϕ*. We then feed the penultimate outputs into the rotamer module for the prediction of side chain conformations, such as *χ*_1_, *χ*_2_, *χ*_3_, *χ*_4_, and *χ*_5_. Lastly, these tensors are used to update the inter-residue module to obtain the final outputs.

**Fig 1.**
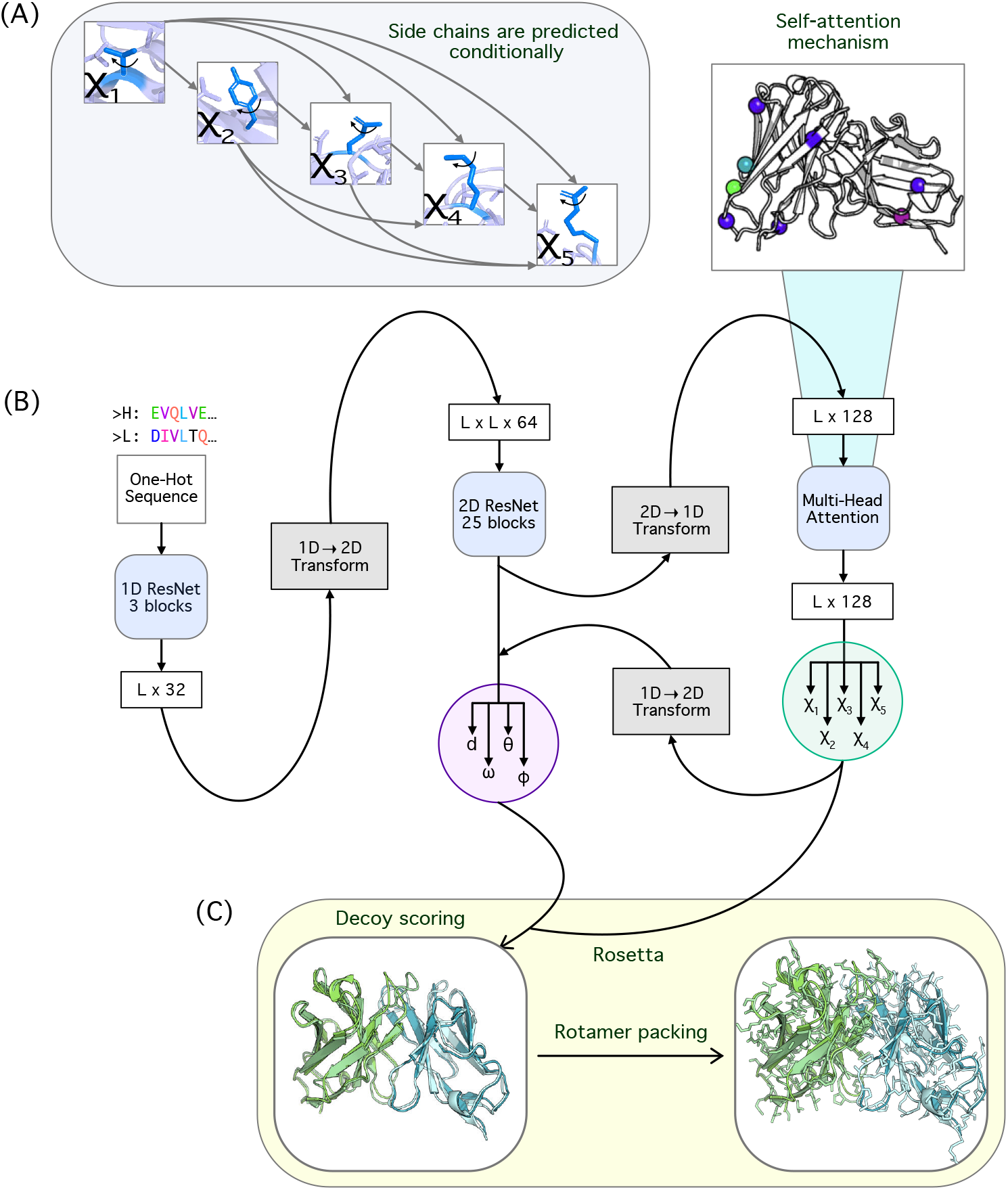
Overview of the DeepSCAb network architecture. (A) Conditional side chain dihedral prediction in DeepSCAb rotamer module with each dihedral after *χ*_1_ depending on previous prediction(s). (B) DeepSCAb architecture for predicting inter-residue geometries and side chain dihedrals. (C) Applications of DeepSCAb for full *F*_V_ realizations and side chain repacking using Rosetta.

We calculate pairwise energies after passing the input sequence through multiple blocks of 1D and 2D ResNets as outlined in the Methods section. The network then splits into four output branches, following the collection of pairwise tensors being fed into the rotamer module. Here, we implemented an attention layer before predicting rotamers. We predicted every *χ* angle after *χ*_1_ by conditioning on preceding *χ* angle(s). For *χ*_1_ angles, we only inputted the inter-residue features.

We additionally experimented with predicting side chains independently using the same training regime (i.e., directly predicting from pairwise features as for *χ*_1_). We observed that predicting side chains conditionally resulted in lower cross-entropy losses for all nine outputs, which is ideal.

### Rotamer module attends to structurally-conserved anchor positions

The rotamer module includes a self-attention layer that allows us to identify the positions that most significantly influence the side chain predictions. Rather than attending broadly across the entire antibody sequence, we observed that the model restricted attention to structurally-conserved residues, which we refer to as anchors.

We tested the conservation of anchor positions in various species and settings including ten human antibody targets, a bovine antibody, and mouse and rat sequences with unknown structures. We collected the human antibodies from the independent test set and selected a random bovine antibody (6E9G). Lastly, the mouse and rat antibody structures shown are predictions from DeepSCAb using the protocol described for DeepAb [14], for random paired sequences from OAS [22]. Across the aforementioned range of systems, we found that the anchor positions, as well as anchor residue types, are frequently conserved (Figure 2A).

**Fig 2.**
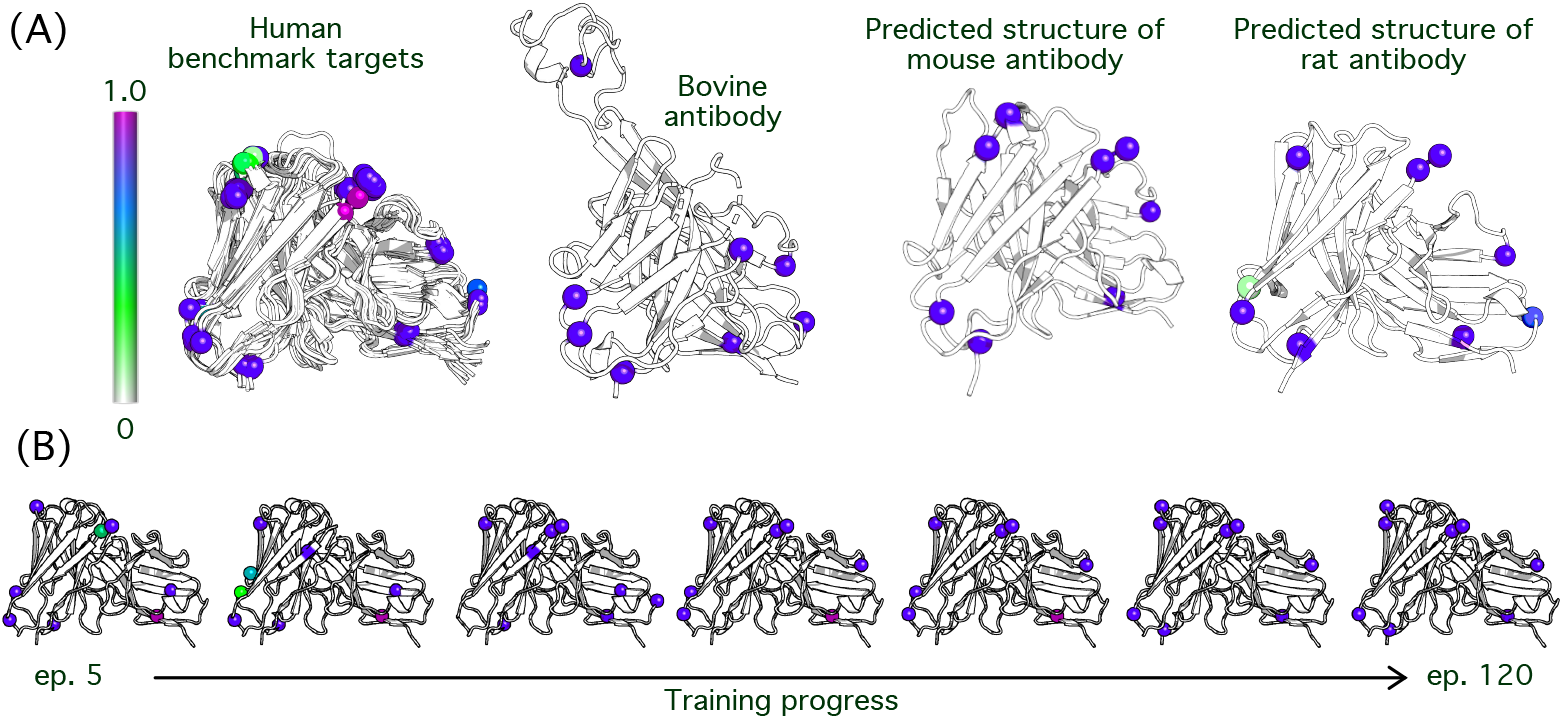
Identification of anchor residue positions from rotamer module attention. Rotamer module attention is interpreted to indicate positional significance in side chain predictions. (A) An attention spectrum (left) ranging from white to magenta represents 0% to 100% attention, respectively. Human, bovine, mouse, and rat antibodies are shown. (B) The variation in attention level is shown with increasing training progress.

When we analyzed how attention patterns changed throughout the course of training we observed a process that resembled a search for anchor residues. The initial epochs (before epoch 25) in training are used as a means of scanning for key positions in the sequence. This subsequently results in switching out a few anchors altogether in the beginning stages of training. The high attention residues begin to settle in their positions at epoch 40, however, the ranges of attention assigned remain dynamic up until late epochs. At epoch 100, the model settles on eight anchor positions commonly with highest levels of attention observed (Figure 2B). The epochs represented in the figure are 5, 20, 40, 60, 80, 100, and 120.

### Side chain predictions improve CDR H3 loop structure accuracy

#### Training on backbone geometries improves side chain predictions

To assess the side chain prediction accuracy of the model without any knowledge of backbone preferences, we designed a control network that consists of primarily the rotamer module. To understand the impact of including backbone conformation for side chain predictions, we evaluated the control network and DeepSCAb on a decoy discrimination task using the RosettaAntibody benchmark. For each target in the benchmark, we score 2800 decoys generated by RosettaAntibody and measure the *RMSD* of the top-1 and top-5 scoring structures referencing the native [23]. For the top-1 scoring decoys, DeepSCAb (*RMSD* = 3.2 Å) outperformed the control network (*RMSD* = 5.0 Å) by 1.8 Å (32 better, 7 same, 10 worse). For the top-5 scoring decoys, DeepSCAb (*RMSD* = 2.5 Å) outperformed the control network (*RMSD* = 3.3 Å) by Å (23 better, 11 same, 15 worse) (Table 1). Due to the considerable improvement observed in DeepSCAb over the control network, we conclude that the inter-residue predictions are necessary for side chain predictions.

**Table 1.** Decoy discrimination compared to DeepSCAb

| Energy | Top 1-Scoring Decoys |  |  |  | Top 5-Scoring Decoys |  |  |  |
| --- | --- | --- | --- | --- | --- | --- | --- | --- |
|  | Better | Same | Worse | *RMSD* | Better | Same | Worse | *RMSD* |
| DeepSCAb | - | - | - | 3.2Å | - | - | - | 2.5Å |
| Control | 32 | 7 | 10 | 5.0Å | 23 | 11 | 15 | 3.3Å |
| DeepH3 | 10 | 33 | 7 | 3.2Å | 10 | 35 | 4 | 2.6Å |

#### Training on side chain geometries improves inter-residual predictions in return

Since DeepSCAb outperformed the control network, we next compared the performance of DeepSCAb to DeepH3 on the same decoy discrimination task. For the Top 1-scoring decoys, DeepSCAb modestly outperformed DeepH3 (10 better, 33 same, 7 worse; Δ*RMSD* = 0 Å). For the Top 5-scoring decoys, DeepSCAb outperformed DeepH3 (10 better, 35 same, 4 worse; Δ*RMSD* = −0.1 Å) (Table 1).

Using the independent test set, we plotted the structures chosen by DeepH3 against the ones chosen by DeepSCAb based on their *RMSD* (Å). DeepSCAb was more successful at distinguishing near-native structures in both Top 1 and Top 5 plots, though improvements over DeepH3 were most notable in the Top 5 comparison (Figure 3A). We then analyzed two targets chosen from the Top 5 decoys for the three methods and *ref2015*. We show the structure scores against *RMSD* in the CDR H3 loop for the target 2FB4 (loop length of 19) (Figure 3B). DeepSCAb outperformed the control network (Δ*RMSD* = −10.8 Å), *ref2015* (Δ*RMSD* = −10.2 Å), and DeepH3 (Δ*RMSD* = −1.2 Å). Comparison of the structures identified by DeepSCAb and DeepH3 revealed that both models are able to place the CDR H3 loop in the correct orientation, however, the addition of side chain information in DeepSCAb results in a more accurate structure (Figure 3C). We further plotted the funnel energies in the CDR H3 loop for the target 3MLR (loop length of 17) (Figure 3D). DeepSCAb outperformed the control network (Δ*RMSD* = −4.4 Å), *ref2015* (Δ*RMSD* = −5.8 Å), and DeepH3 (Δ*RMSD* = −1.4 Å). We displayed all structures and the native overlaid for the target 3MLR (Figure 3E). DeepSCAb predicts the loop structure with the highest accuracy, given one of the longer and more difficult of the CDR H3 loops. Hence, the addition of side chain orientations is beneficial for accurately predicting pairwise geometries.

**Fig 3.**
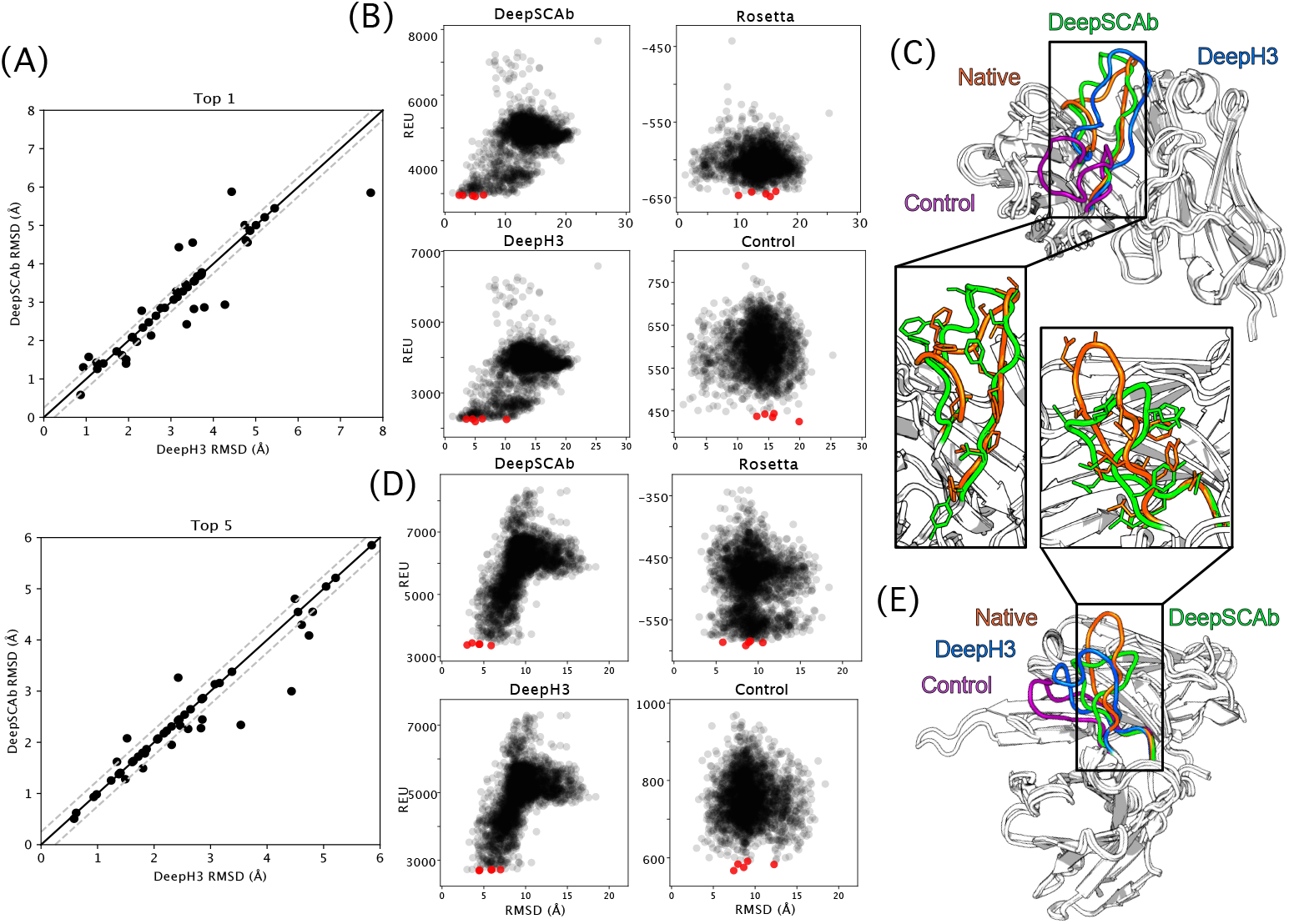
Comparison of CDR H3 structure prediction accuracy. The accuracy with which DeepSCAb, DeepH3, and the control network predict the CDR H3 loop structure is measured via decoy structure scoring tasks. (A) In Top 1-scoring decoy structures (top) and Top 5-scoring structures (bottom), the performance of DeepSCAb is compared to DeepH3 using the test set. (B) The CDR H3 energies for the three methods and Rosetta (ref2015) that correspond to 2FB4 are plotted against their *RMSD*. The five best scoring structures for each plot are indicated in red. (C) The best prediction from Top 5-scoring decoys for target 2FB4 are shown for DeepSCAb (green, 2.34 Å *RMSD*), DeepH3 (blue, 3.535 Å *RMSD*), and the control network (purple, 13.091 Å *RMSD*) all compared to the native (orange). (D) The CDR H3 energies for the three methods and Rosetta (ref2015) that correspond to 3MLR are plotted against their *RMSD*. The five best scoring structures for each plot are indicated in red. (E) The best prediction from Top 5-scoring decoys for target 3MLR are shown for DeepSCAb (green, 2.998 Å *RMSD*), DeepH3 (blue, 4.432 Å *RMSD*), and the control network (purple, 7.441 Å *RMSD*) all compared to the native (orange).

### DeepSCAb is competitive with alternative rotamer packing methods

The context of the predicted side chains is crucial in determining the accuracy and usefulness of a method. Side chains that are greatly exposed to a solvent play an active role in the binding of an antigen, yet are also inherently the most flexible. We evaluated the performance of our method and three alternative methods as a function of relative side chain solvent accessible surface area (SC SASA) using Rosetta. We compared the success of DeepSCAb in predicting side chain conformations to PEARS, SCWRL4, and Rosetta, where the native structure was used as a reference for all measurements. We omitted the target 3MLR from side chain packing and relative SC SASA comparisons as PEARS was unable to model this structure due to its long L3 loop.

In Figure 4, we illustrate that while DeepSCAb did not significantly outperform any method in the *F*_V_ region, the CDR H3 loop or the non-H3 loops, it remained competitive in per-residue side chain prediction accuracy with the most successful alternative method, PEARS. However, we note that the alternative methods have and require access to the true backbone, while our method does not. We took a closer look at the average *RMSD* at various relative SC SASA values and found that DeepSCAb was able to uncover similar amounts of information from only the sequence, even for the challenging CDR H3 loop. As the methods, on average, perform very similar to one another (Table 1), DeepSCAb may be the most useful in antibody engineering applications, where the backbone conformation is flexible upon design requirements and restrictions.

**Fig 4.**
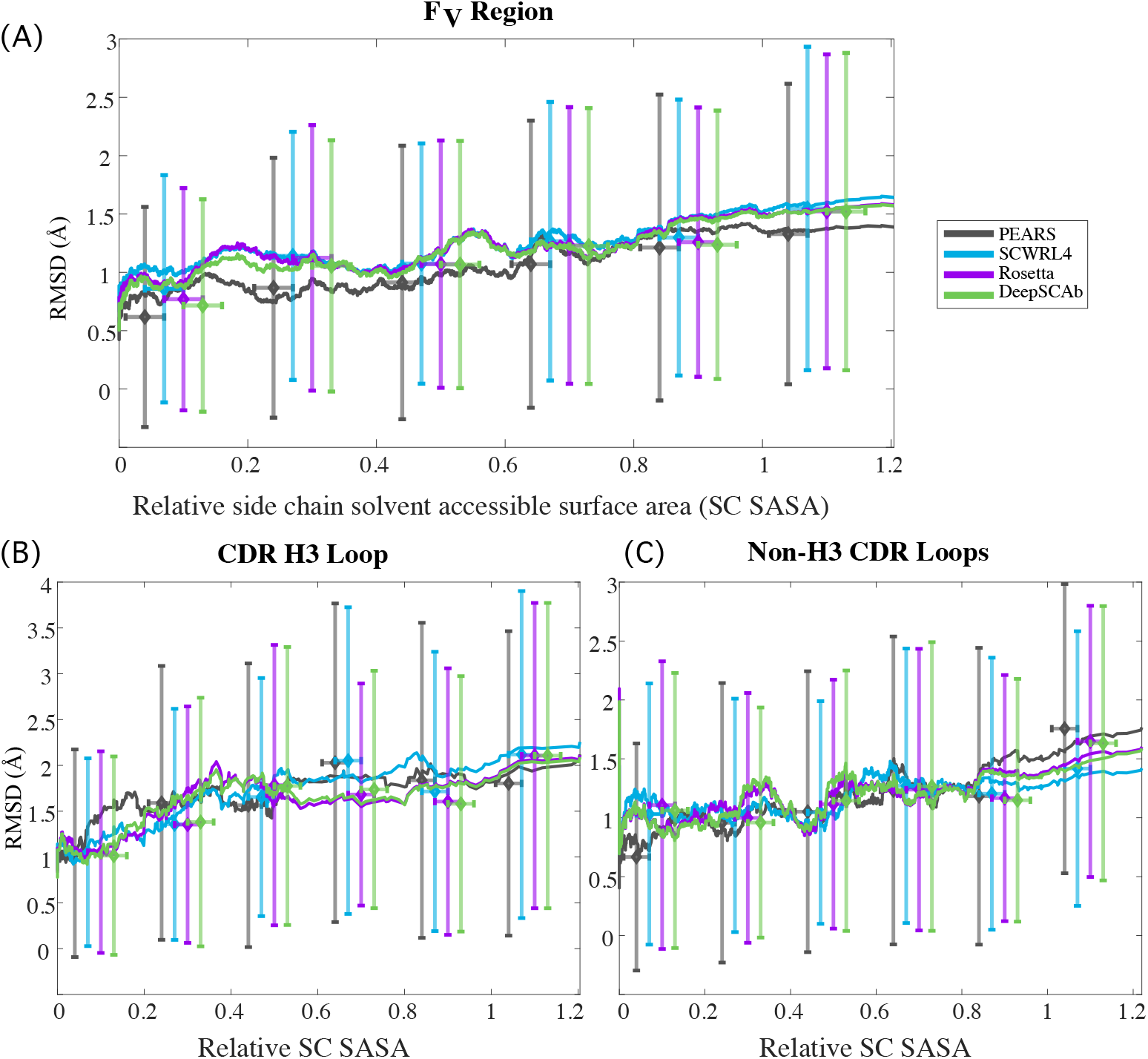
Comparison of side chain prediction accuracy with increasing solvent exposure. Plots show moving means of side chain *RMSD* as a function of relative side chain solvent accessible surface area (SC SASA), with error bars indicating standard deviation within each 0.2 interval and horizontal ticks corresponding to mean *RMSD*. (A) Comparison of *F*_V_ side chain prediction based on per-residue *RMSD* as a function of SC SASA for PEARS, SCWRL4, Rosetta, and DeepSCAb, all referencing the native structure, with moving average window length of 400 datapoints. (B) Comparison of CDR H3 loop side chain prediction accuracy with increasing relative SC SASA (window length of 60 datapoints). (C) Comparison of non-H3 CDR loops’ combined side chain prediction accuracy with increasing relative SC SASA (window length of 120 datapoints).

## Discussion

The results outlined in our work show that our method is a step towards accurate antibody docking and design via inclusion of side chain predictions. We demonstrated that DeepSCAb predictions remain competitively accurate at varying side chain surface exposure. Compared to alternative rotamer building methods, DeepSCAb is robust to non-canonical loops as it learns patterns pertaining to neighbor backbone, side chain geometry, and sequence, rather than relying on rotamer libraries. Using the rotamer module attention, we are able to identify the residue types and positions that are the most influential in the context of side chain predictions. While this analysis provides insight into rotamer prediction, it does not reveal local biophysical interactions that could be tied into the fundamentals of side chain conformation in complex energy landscapes. The anchor sites are consistently scattered throughout the sequence, in stark contrast with the local chemical environment typically considered by most side chain placement algorithms. Perhaps DeepSCAb is learning an internal, structurally conserved numbering scheme as a reference for side chain prediction similar to the well-performing PEARS algorithm [18], which was designed intentionally to use antibody-specific residue positioning to condition rotamer predictions. Alternatively, the model could be identifying global antibody features such as germline class or species, which have some side chains conserved.

This work demonstrated that inter-residue features improve side chain predictions. Also, inclusion of side chains improves overall *F*_V_ structure prediction compared to machine learning models that only predict inter-residue geometries. Concurrent with this work, improved methods for antibody structure prediction have been developed [14, 15]. DeepAb uses a similar architecture to predict inter-residue geometries, and ABlooper predicts CDR loop coordinates directly. Our work suggests that both of these methods might be improved by incorporating side chain context into predictions.

Most side chain repacking methods average sample conformations based on a backbone-dependent rotamer library [17], and accurate methods for antibody side chain repacking estimate *χ* angle densities based on a position-dependent rotamer library [18]. Since our method does not require structure as an input, DeepSCAb is unaffected by uncertainty in backbone structure [24] and allows for predictions on sequences without known backbone conformations. This feature is useful when there are multiple potential backbone conformations of interest, e.g., for the design of new therapeutic antibodies. With minimal modification, our network can aid antibody design. For instance, DeepSCAb can be used in parallel with RosettaAntibodyDesign [25] for rapid placement of side chains or to hallucinate new antibody sequences using the trRosetta architecture [10].

## Conclusion

In this study, we investigated the effect of inter-residual predictions on the accuracy of side chain dihedrals as well as the effect of rotamer predictions on the overall antibody structure prediction accuracy. We found that DeepSCAb competitively predicts rotamers when compared to alternative methods that require true backbone coordinates. The performance of our method is robust to when the backbone is perturbed or deviates from the crystal structure. Since DeepSCAb predicts a probability distribution over the backbone and side chain geometries, we expect it will be adaptable to and useful for designing new antibodies.

## Acknowledgments

This work was supported by National Science Foundation Research Experience for Undergraduates grant DBI-1659649 (D.A.), AstraZeneca (J.A.R.), National Institutes of Health grants T32-GM008403 (J.A.R.) and R01-GM078221(S.P.M., J.J.G.). Computational resources were provided by the Maryland Advanced Research Computing Cluster (MARCC).

Dr. Gray is an unpaid board member of the Rosetta Commons. Under institutional participation agreements between the University of Washington, acting on behalf of the Rosetta Commons, Johns Hopkins University may be entitled to a portion of revenue received on licensing Rosetta software including methods discussed/developed in this study. As a member of the Scientific Advisory Board, J.J.G. has a financial interest in Cyrus Biotechnology. Cyrus Biotechnology distributes the Rosetta software, which may include methods developed in this study. These arrangements have been reviewed and approved by the Johns Hopkins University in accordance with its conflict-of-interest policies.

## Supporting information

**Supp. Table 1.** Average *RMSD* (in Å) at relative SC SASA values for the CDR H3 loop

| Method | Relative SC SASA values |  |  |  |  |  |
| --- | --- | --- | --- | --- | --- | --- |
|  | 0.1 | 0.3 | 0.5 | 0.7 | 0.9 | 1.1 |
| PEARS | 1.041 | 1.591 | **1.565** | 2.029 | 1.837 | **1.803** |
| SCWRL4 | 1.052 | 1.357 | 1.654 | 2.052 | 1.716 | 2.118 |
| Rosetta | 1.053 | **1.353** | 1.785 | **1.682** | 1.605 | 2.107 |
| DeepSCAb | **1.015** | 1.382 | 1.775 | 1.737 | **1.580** | 2.107 |

**Supp. Figure 1.**
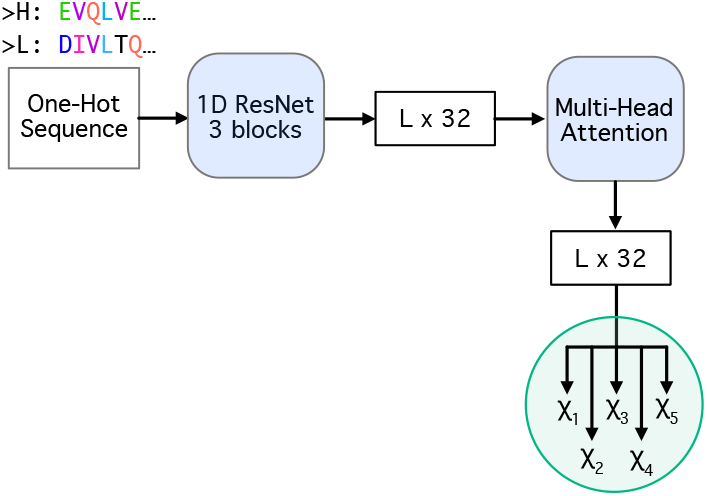
Side chain-only control network architecture. The control network has a similar architecture to the rotamer module in DeepSCAb.

**Supp. Figure 2.**
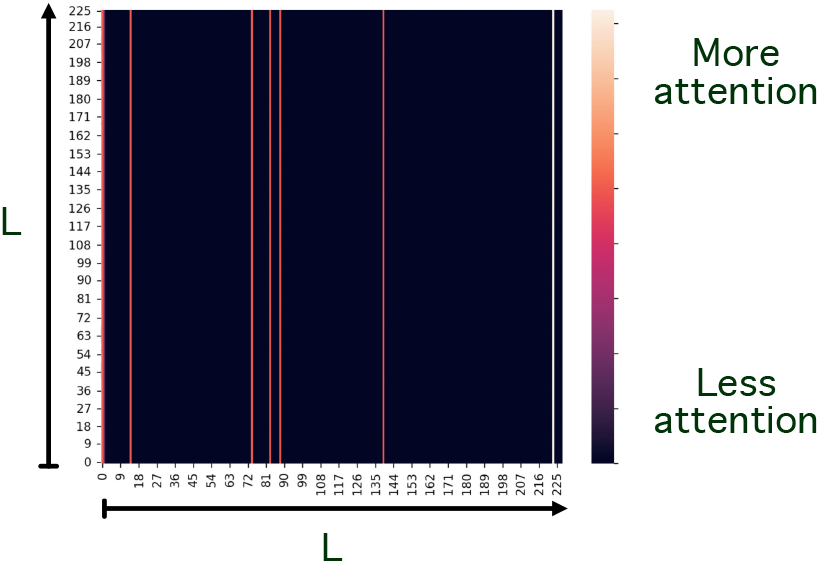
Interpretation of the attention matrices. *L* × *L* matrix is shown for a target chosen at random. The final attention is calculated by averaging over the rows to collapse the matrix to dimension *L* × 1.

**Supp. Figure 3.**
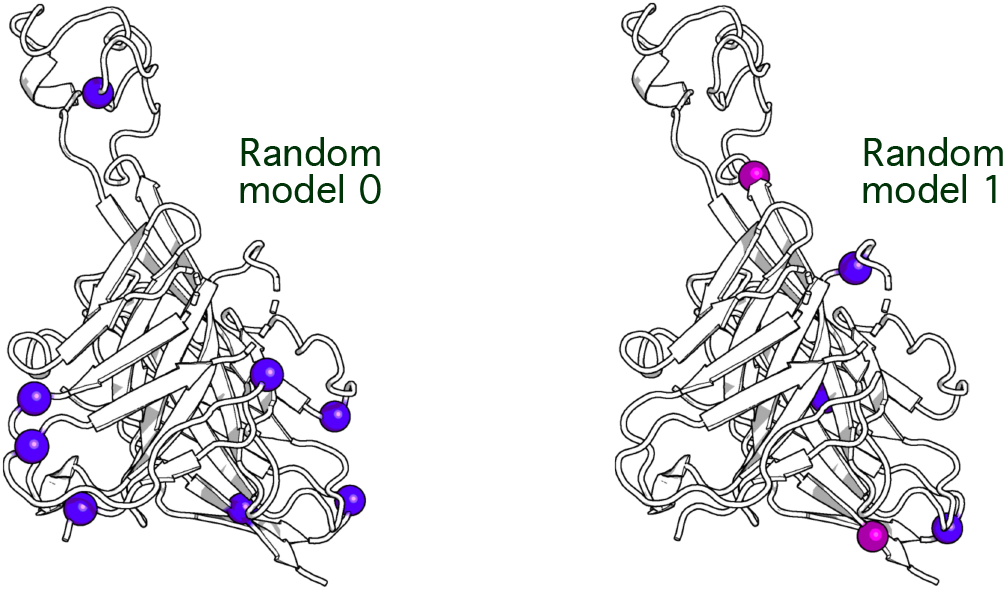
Comparison of anchor positions between two random models of DeepSCAb. The bovine PDB 6E9G is interpreted by two different models to show the variation of anchor positions.

## References

1. Sela-Culang I, Kunik V, Ofran Y. The structural basis of antibody-antigen recognition. Frontiers in Immunology. 2013;4:1–13.

2. Tsuchiya Y, Mizuguchi K. The diversity of H3 loops determines the antigen-binding tendencies of antibody CDR loops. Protein Science. 2016;4(25):815–825.

3. Leem J, Dunbar J, Georges G, Shi J, Deane CM. ABodyBuilder:Automated antibody structure prediction with data-driven accuracy estimation. mAbs. 2016;7(8):1259–1268.

4. Schritt D, Li S, Rozewicki J, Katoh K, Yamashita K, Volkmuth W, Cavet G, Standley DM. Repertoire Builder: High-throughput structural modeling of B and T cell receptors. Molecular Systems Design and Engineering. 2019;4(4):761–768.

5. Weitzner BD, Jeliazkov JR, Lyskov S, Marze N, Kuroda D, Frick R, Adolf-Bryfogle J, Biswas N, Dunbrack RL, Gray JJ. Modeling and dicking of antibody structures with Rosetta. Nature Protocols. 2017;2(12):401–416.

6. Spassov V, Yan L, Flook P. The dominant role of side-chain backbone interactions in structural realization of amino acid code. ChiRotor: A side-chain prediction algorithm based on side-chain backbone interactions. Bioinformatics. 2007.

7. Chiu ML, Goulet DR, Teplyakov, Gilliland GL. Antibody Structure and Function: The Basis for Engineering Therapeutics. Antibodies. 2019;8(4):55.

8. Reichert JM. Antibodies to watch in 2017. mAbs. 2017;9(2):167–181.

9. Gao W, Mahajan SP, Sulam J, Gray JJ. Deep Learning in Protein Structural Modeling and Design. Patterns. 2020;1(9):100–142.

10. Yang J, Anishchenko I, Park H, Peng Z, Ovchinnikov S, Baker D. Improved protein structure prediction using predicted interresidue orientations. Proceedings of the National Academy of Sciences of the United States of America. 2020;3(117):1496–1503.

11. Jumper J, Evans R, Pritzel A, Green T, Figurnov M, Ronneberger O, Tunyasuvunakool K, Bates R, Žídek A, Potapenko A, Bridgland A, Meyer C, Kohl SAA, Ballard AJ, Cowie A, Romera-Paredes B, Nikolov S, Jain R, Adler J, Back T, Petersen S, Reiman D, Clancy E, Zielinski M, Steinegger M, Pacholska M, Berghammer T, Bodenstein S, Silver D, Vinyals O, Senior AW, Kavukcuoglu K, Kohli P, Hassabis D. Highly accurate protein structure prediction with AlphaFold. Nature. 2021.

12. Baek M, DiMaio F, Anishchenko I, Dauparas J, Ovchinnikov S, Lee GR, Wang J, Cong Q, Kinch LN, Schaeffer RD, Millán C, Park H, Adams C, Glassman CR, DeGiovanni A, Pereira JH, Rodrigues AV, van Dijk AA, Ebrecht AC, Opperman DJ, Sagmeister T, Buhlheller C, Pavkov-Keller T, Rathinaswamy MK, Dalwadi U, Yip CK, Burke JE, Garcia KC, Grishin NV, Adams PD, Read RJ, Baker D. Accurate prediction of protein structures and interactions using a three-track neural network. Science. 2021.

13. Ruffolo JA, Guerra C, Mahajan SP, Gray JJ. Geometric potentials from deep learning improve prediction of CDR H3 loop structures. Bioinformatics. 2020.

14. Ruffolo JA, Sulam J, Gray JJ. Antibody structure prediction using interpretable deep learning. 2021.

15. Abanades B, Georges G, Bujotzek A, Deane CM. ABlooper: Fast accurate antibody CDR loop structure prediction with accuracy estimation. 2021.

16. Cohen T, Halfon M, Schneidman-Duhovny D. NanoNet: Rapid end-to-end nanobody modeling by deep learning at sub angstrom resolution. 2021.

17. Krivov GG, Shapovalov MV, Dunbrack RL. Improved prediction of protein side-chain conformations with SCWRL4. Proteins: Structure, Function and Bioinformatics. 2009;4(77):778–795.

18. Leem J, Georges G, Shi J, Deane CM. Antibody side chain conformations are position-dependent. Proteins: Structure, Function and Bioinformatics. 2018;4(86):383–392.

19. Dunbar J, Krawczyk K, Leem J, Baker T, Fuchs A, Georges G, Shi J, Deane CM. SAbDab: The structural antibody database. Nucleic Acids Research. 2014;D1(42):1140–1146.

20. Adolf-Bryfogle J, Xu Q, North B, Lehmann A, Dunbrack RL. PyIgClassify: a database of antibody CDR structural classifications. Nucleic acids research. 2014;43(D1):D432–D438.

21. Vaswani A, Shazeer N, Parmar N, Uszkoreit J, Jones L, Gomez AN, Kaiser L, Polosukhin I. Attention Is All You Need. 31st Conference on Neural Information Processing Systems. 2017.

22. Kovaltsuk A, Leem J, Kelm S, Snowden J, Deane CM, Krawczyk K. Observed Antibody Space: A Resource for Data Mining Next-Generation Sequencing of Antibody Repertoires. The Journal of Immunology. 2018;8(201):2502–2509.

23. Jeliazkov J, Frick R, Zhou J, Gray JJ. Robustification of RosettaAntibody and Rosetta SnugDock. PLoS ONE. 2021;3(16):1–20.

24. Schwarz D, Georges G, Kelm S, Shi J, Vangone A, Deane CM. Co-evolutionary distance predictions contain flexibility information. Bioinformatics. 2021.

25. Adolf-Bryfogle J, Kalyuzhniy O, Kubitz M, Weitzner BD, Hu X, Adachi Y, Schief WR, Dunbrack RL. RosettaAntibodyDesign (RAbD): A General Framework for Computational Antibody Design PLoS Computational Biology. 2018;4(14).

